# ^13^C metabolic flux ratio analysis by direct measurement of free metabolic intermediates in L. mexicana using gas chromatography-mass spectrometry

**DOI:** 10.1101/073676

**Authors:** Milica Ng, Eleanor C. Saunders, Moshe Olshansky, Malcolm J. McConville, Vladimir A. Likić

## Abstract

Until recently, ^13^C-based flux analyses have almost exclusively relied on analysis of labelled amino acids in proteins. This approach is not directly applicable to *Leishmania*, as these parasites scavenge most of their amino acids from the media. *Leishmania* are also unusual in that they i) share little genomic similarity with other organisms ii) constitutively express their metabolic genes and iii) display minimal changes in the enzyme levels throughout their life cycle stages. The three factors have contributed to an early development of comprehensive and reproducible ^13^C-based metabolomics approaches in these parasites. The work presented here contributes to the creation of new ^13^C-based metabolic flux approaches based on the isotopologue analysis of free metabolite pools in *Leishmania mexicana*. Namely, a new approach is presented for simultaneous calculation of *in vivo* fractional fluxes (or flux ratios) into two or more metabolite nodes with carbon dioxide condensation, based on isotopologue analysis of free metabolite pools. This method is used to perform the first quantitative *in vivo* fractional flux calculation of central carbon metabolism in any human parasite.

## Introduction

Current metabolic research focuses on how to harness the newly acquired omics data in order to be able to i) restore metabolic function when things go wrong (e.g. in numerous metabolic diseases), ii) disable metabolism of pathogens, and iii) optimise metabolic production rates of biotechnologically relevant organisms. Most importantly we want to be able to alter cellular metabolism in a predictable, rather than previously ad hoc, manner. The cellular metabolism comprises a large number of biochemical reactions, and the reconstruction of the metabolic network topology is an important first step in understanding metabolic function in a particular cell. This static picture gives information on the cell’s metabolic capacity. What is of more interest however, is the amount of activity throughout different parts of the network. Metabolic flux, defined as the rate of flow of metabolite atoms through the metabolic network, is the direct measure of a metabolic pathway’s (or individual reaction’s) activity level.

In general, metabolic flux inside the cell is regulated by altering the number and/or the capacity of enzymes that catalyse a particular reaction. Therefore flux increase or decrease is often hypothesized based on the measurements of the relevant enzyme or metabolite levels. This can often lead to incorrect conclusions, as an increase in enzyme levels does not necessarily indicate an increase in the metabolic flux, as enzymes can be present but inactive [1,2]. Alternatively, an enzyme can be active without having any significant control of the overall pathway flux (this scenario is the subject of metabolic control analysis [3]). Similarly, metabolite levels alone are not accurate indicators of enzymatic activity. The tendency of an organism, or a cell, to maintain internal equilibrium by adjusting its physiological processes (homeostasis) often results in metabolite pool levels remaining constant even though a coordinated increase, or decrease, of enzymatic activity throughout a pathway is taking place [4]. Moreover, metabolite levels can change in the opposite direction to the related enzyme’s activity, if enzymes upstream or downstream of them have been activated or deactivated. Therefore, data on enzyme and metabolite levels are only able to generate predictions about the rate of flow of metabolite atoms between different nodes. Metabolic flux on the other hand, is the true measure of this rate.

Only a small number of fluxes, such as nutrient uptake, product secretion and biomass production can be measured directly. The intracellular fluxes, which carry the most important information, are not directly measurable. Different mathematical modelling strategies have been developed to solve the problem. According to whether they measure fluxes as a function of time or constant fluxes at metabolic steady state, they can be divided into dynamic and static (or steady-state) modelling approaches respectively. This work is focused on the latter.

All living cells maintain steady state by regulatory mechanisms in order to compensate for changes in their external environments, e.g. temperature, nutrient content or pH level. It has been observed that even after relatively large environmental perturbations, cells reach a new steady state within minutes. Measuring fluxes in metabolic equilibrium, reached after perturbations of interest, provides valuable information on the adjustments that cells make to regulate metabolic function. At metabolic steady state, metabolite levels in the network are constant (or assumed to be near enough constant compared to the relevant flux values). Thus, based on the law of mass conservation, the sum of all the fluxes that add to a particular intracellular metabolite pool must equal the sum of all the fluxes that deplete the same pool. Therefore, utilising known reaction stoichiometry, a set of linear equations can be written for each intracellular metabolite pool inside the metabolic network. The number of equations in the model will be equal to the number of metabolites in the network, and the number of unknowns will be equal to the number of reactions (or flux values to be determined). In a typical metabolic network, metabolites are shared between reactions and hence the number of metabolites will be smaller than the number of reactions. This means that the overall linear equation system will be underdetermined. In other words, there will be more unknowns than the equations to solve for them. Two different strategies have been devised to deal with this: i) flux balance analysis [5,6,7] and ii) ^13^C-based flux analysis. Our focus is on the latter.

The two most prominent ^13^C-based approaches are: ^13^C metabolic flux analysis (^13^C-MFA) [8,9,10,11], and ^13^C metabolic flux ratio analysis (^13^C-MFRA) [12,13,14]. Typically, ^13^C-labelled substrates are fed to the cells under study, allowing different metabolic pathways to cleave and scramble substrates' carbon backbone. Under the conditions of metabolic and isotopic steady-state, isotopic composition of metabolic intermediates depends on the labelling state of the input substrates, network topology, and fluxes operating in the network. The isotopic composition of intermediates (and products) can be measured by NMR [15,16,17] or mass spectrometry [18,19,20], and given that the input substrates and the network topology are known, fluxes operating in the metabolic network can be inferred by mathematical modelling. However, underlying this is a significant mathematical modelling effort, and a number of challenges remain. Firstly, ^13^C-MFA requires that the topology of the metabolic network is known, i.e., as a part of mathematical modelling the network topology is assumed. This in turn implies that it is possible to assume incorrect network topology, and obtain a seemingly reasonable solution (fluxes that fit the experimental data), without any warning signs. Secondly, because of the integral nature of the model, any errors in the input data will propagate to affect the entire solution. Once the solution fluxes are obtained by ^13^C-MFA a detailed statistical evaluation of the results is recommended [21].

^13^C metabolic flux ratio analysis (^13^C-MFRA) is a ‘divide and conquer’ variation on ^13^C-MFA. Namely, the method relies on a set of measurements from ^13^C-labelling experiments, at isotopic steady state, to resolve relative flux contributions of metabolic pathways and/or reactions to a single metabolic network node of interest. Unlike ^13^C-MFA, it does not require comprehensive network topology. This comes at the price of being able to compute only relative and not net flux values. However, in organisms where topology is yet unknown this strategy can prove invaluable in deciphering topology before embarking on a more elaborate ^13^C-MFA work. The widely available detailed drawings of metabolic networks can often leave a novice with a belief that very little, if anything, is unknown due to the evolutionary preservation of metabolic pathways. However, a recent review estimates that 30-40% of metabolic activities are missing enzyme and/or gene allocation in an average organism [22]. The view of enzymes as a single substrate converting catalyst is also challenged by the existence of so called promiscuous enzymes which have the ability to catalyse more than one reaction [23,24]. Therefore, although often unjustifiably neglected, ^13^C-MFRA offers an important insight into *in vivo* flux activity around individual network nodes. This approach is also essential to reduce the overall computational effort when the biological question of interest can be answered by considering only a few metabolic nodes.

*Leishmania* cause the human disease leishmaniasis, the clinical symptoms of which range from cutaneous and mucocutaneous forms through to visceral leishmaniasis that is fatal unless treated. It is estimated that 12 million people are currently infected worldwide, and that more than 350 million people are at risk [25,26]. The leishmaniasis is thought to result in 60,000 deaths each year [27], making it the second deadliest protozoan infection after malaria. There is no vaccine against leishmaniasis [28] and current chemotherapies are often costly, cause severe side effects, and demonstrate increasing rates of resistance [29,30,31]. There is an urgent need to identify new drug targets, with metabolic pathways that are necessary for growth, differentiation and virulence of these important human pathogens being likely candidates. These pathways must, at the same time, not present or at least be minimally utilised by their host. As well as targeting these novel parasite-specific pathways, it may be possible to create metabolic ‘bottlenecks’ leading to the accumulation of toxic intermediates. Alternatively, inhibition of an enzyme with slow turnover in a parasite and fast turnover in the host is an option (e.g. [32]). Studying mechanisms by which these pathogens acquire drug resistance is another important area of research.

In microorganisms, ^13^C-based flux analyses have almost exclusively relied on analysis of labelled amino acids isolated from hydrolysed proteins. In prokaryotes, this approach is favoured as these organisms synthesise many of their amino acids *de novo*. Even though incorporation of ^13^C atoms into protein takes much longer then into the cellular precursors (i.e. proteinogenic amino acids), the abundance of protein within the cell makes it an easy substrate to isolate and analyse. Furthermore, in ^13^C-based flux analyses, out of 20 amino acids, 8 metabolite patterns are usually discernible and this is often enough to constrain the stoichiometric model of central carbon metabolism [33]. Furthermore, metabolite extraction from bacteria is still an unsolved problem [33]. These analyses, however, are hampered in organisms that scavenge extracellular amino acids, such as *Leishmania spp*. For example *L. mexicana* have been shown to scavenge most of their amino acids from the media and, with the exception of alanine, glutamate and aspartate, the labelling of other amino acids is negligible [34].

In common with analyses in other eukaryotes, accurately determining flux in *Leishmania spp.* is further complicated by unknown gene functionality and network topology as well as the intracellular compartmentalisation of metabolites and pathways. For example, the GeneDB database indicates that 60% of *L. major* genes currently have no known functionality [35,36]. The knowledge of metabolic gene function is necessary, but not sufficient. The connectivity and arrangement of metabolic pathways need to be experimentally elucidated for each organism of interest. Nonetheless some progress has been made in identifying important carbon sources and delineating the network topology of central carbon metabolism [34]. This was made possible by using a range of ^13^C-substrates and detecting the ^13^C-enrichment of metabolic intermediates. Despite these advances, ^13^C-based flux analysis has not been applied to *Leishmania spp*.

The study of flux throughout central carbon metabolism in *Leishmania spp.* is of particular interest given the pathways’ central roles in parasite growth and virulence [37]. This is despite the observation that central carbon metabolism is largely conserved between species as it is anticipated that, given the significant differences in compartmentalisation and regulatory mechanisms between host and parasite, central carbon metabolism remains a good drug target in *Leishmania spp*. ^13^C-MFRA is attractive means to delineate central carbon metabolism flux in *Leishmania spp*. Firstly, ^13^C-MFRA requires only the knowledge of reactions which top up a metabolite pool of interest and, as a result, the calculation is often largely independent of the overall network topology. This is particularly useful given methods to experimentally ascertain *Leishmania*’s metabolic network topology are still evolving. Secondly, ^13^C-MFRA is amenable to high-throughput analysis, and coupled with the advances in GC-MS techniques, can lead to the development of rapid methods capable of scanning a multitude of different experimental set ups within reasonable time frames.

We present here a novel approach for simultaneous calculation of *in vivo* fractional fluxes (or flux ratios) into two or more metabolite nodes with carbon dioxide condensation, based on isotopologue analysis of free metabolite pools. This method highlights the yet unexplored potential of combining ^13^C-MFRA and direct metabolomics measurements for the quantitative study of metabolism. In particular the approach will benefit the study of i) the organisms whose metabolic network topology is poorly understood, and ii) the organisms in which a sufficient number of metabolite mass isotopic distributions cannot be inferred from the measurements of proteinogenic amino acids. The utility of the approach is demonstrated by its application on the data collected for the malate and oxaloacetate nodes in central carbon metabolism of rapidly dividing *L. mexicana* parasites, providing the first in vivo quantitative fractional flux values for *Leishmania spp.*, or any other protozoan. These are also, to the best of our knowledge, the first fractional flux results based on the mass isotopic distribution measurements of free intracellular metabolic intermediates.

## Methods

### Notation

In this section we introduce the notation used throughout. The property that will be of central interest in considerations below is mass isotopic distribution vector **I**. Let **s** denote the vector of a single chemical species that differ in their isotopic composition s = [s_0_, s_1_, …, s_n_] (here chemical species can be either a single atom or a molecule). Let **m** denote the vector of corresponding masses **m** = [m_0_, m_1_, …, m_n_], such that m_0_ is the mass of the isotopic species s_0_, m_1_ is the mass of the isotopic species s_1_, and so on. Let **I** denote the isotopic distribution vector **I** = [a_0_, a_1_, …, a_n_] where the numbers a_0_, a_1_ … a_n_ represent abundances of individual isotopic species with masses m_0_, m_1_, …, m_n_, such that a_0_ + a_2_ + … + a_n_ = 1. In the standard convention the vectors **s**, **m** and **I** are ordered according to the increasing mass (m_0_ < m_1_ < … < m_n_) such that a_0_ is the abundance of the isotopic species s_0_ which has the smallest mass m_0_. Mass spectrometry (MS) measures *m/z* ratios to produce intensity patterns called mass spectra. The mass spectrum observed experimentally by MS, is directly related to the mass isotopic distribution of the chemical species under observation. In case of singly charged species (z=1) MS measures directly atomic/molecular masses of the species involved. Most mass spectrometers can distinguish masses of 0.05 u apart, however detected masses are commonly binned in integer increments. In metabolomics, all masses detected in the range from −0.3 to +0.7 relative to an integer mass are usually summed and recorded as the integer mass intensity.

There are two distinct ways in which two isotopic populations of atoms can combine into a complex mixture: simple mixing and chemical bonding. In simple mixing of two chemical species with well-defined isotopic populations **I**_1_ and **I**_2_, the mass isotopic distribution of the resulting population is given by:

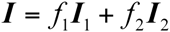

where f_1_ and f_2_ are mixing fractions of the two populations.

In mixing due to chemical bonding (i.e. mixing of isotopic species occurs due to chemical reactions) atoms combine in fixed stoichiometric ratios. If the two populations of chemical species are characterized by mass isotopic distributions **I**_1_ and **I**_2_, the resulting chemical species will have the mass isotopic distribution:

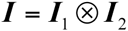

where '⊗' denotes discrete convolution (also known as Cauchy product [38]). This simple relationship is a direct result of integer binning of detected masses.

### A metabolite node without condensation or cleavage

Consider two metabolic pathways converging to a single metabolite within a larger metabolic network. Figure 1A shows two metabolites, D and E, both converted to G. Let us assume that isotopic populations of D and E are well defined, and given by **I**_D_ and **I**_E_ respectively. If no condensation or cleavage occurs in either of the two reactions D → G and E → G, then G inherits the carbon backbone directly from D or E. If the metabolic network is in metabolic and isotopic steady state, the resulting mass isotopic distribution of G will be a result of simple mixing, given as a combination of the two mass isotopic distributions in proportions determined by relative contributions of fluxes v_D_ and v_E_:

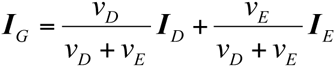

or

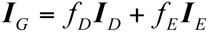

where f_D_ and f_E_ are fractional contributions of D → G and E → G reactions, respectively. The above equation can be rewritten in a matrix form:

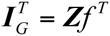

where **f** = [f_D_, f_E_] contains mixing fractions, and f_D_ + f_E_ = 1. The matrix **Z** contains mass isotopic distributions and its columns are mass isotopic distributions for individual chemical species arranged in the same order as the fractional contributions in the vector f (i.e. 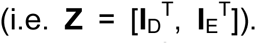). If one is able to measure **I**_D_, I_E_, and **I**_G_, the fractional contributions of the two reactions f_D_ and f_E_ can be obtained by solving the above matrix equation for **f**. In general, **Z** is not a square matrix. However, in the majority of central carbon metabolism reactions which are of interest in the labeling experiments, the number of distinct isotopic populations being mixed is at most 3 (number of **Z** columns), and the smallest number of carbon atoms in their skeletons is at least 3, hence the number of **Z** rows is at least 4. Therefore, the number of rows will be greater than the number of columns. Hence, provided that isotopic distribution vectors of the populations entering the node are linearly independent, **Z** will have a left inverse:

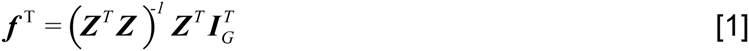

**Figure 1:**
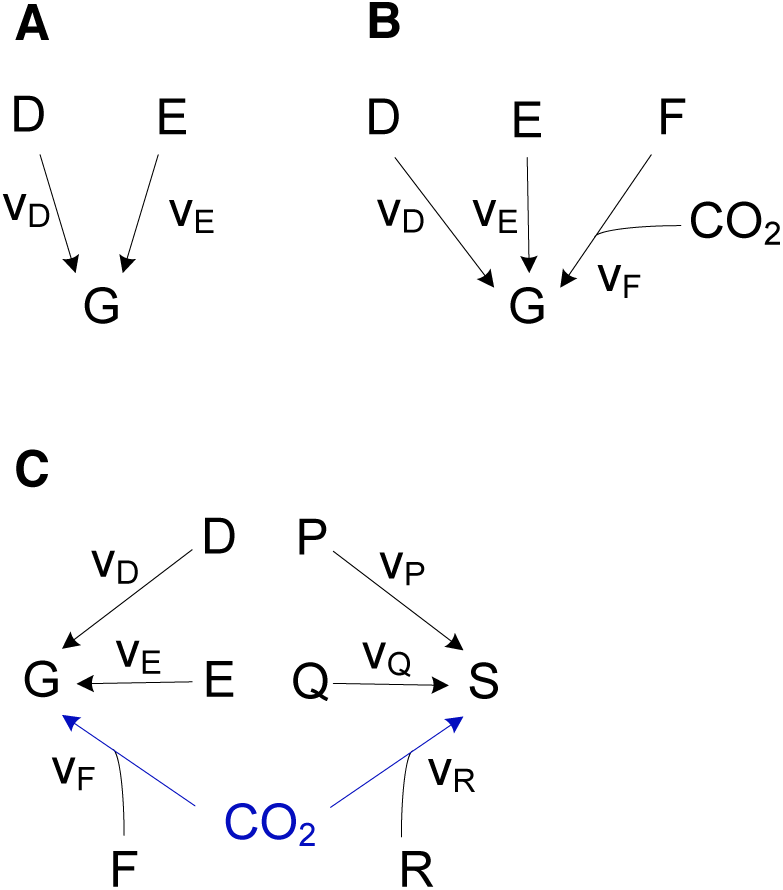
Three scenarios for converging metabolites. Panel (A): metabolites D and E are converted to metabolite G without condensation or cleavage. In this case G inherits the carbon backbone directly from both D and E. Panel (B): three metabolites D, E, and F contribute carbon backbones to metabolite G. In this case metabolites D and E carbon backbones are form metabolite’s G backbone without condensation or cleavage, while the metabolite F’s backbone combines with the carbon in CO_2_ to form the carbon skeleton of G. Panel (C): similar to (B) except that there are now two nodes coupled via CO_2_

We note that the Equation 1 can be extended to a node where k metabolites converge without condensation or cleavage, with mass isotopic distributions **I**_D_, I_E_, …, **I**_M_, and respective fractional contributions vector is **f** = [f_D_, f_E_, …, f_M_], provided that left inverse of **Z** exists.

### A metabolite node with CO_2_ condensation

The situation where two or more metabolites converge to a single node without condensation or cleavage is rare. Consider a prototypical model of the TCA cycle with several anaplerotic reactions, as shown in Figure 2. In this schematic, five metabolite nodes can be identified as candidates for flux ratio determination: PEP, Pyr, OAA, Mal, and Suc (on the basis that these metabolites’ carbon backbones are produced in more than one reaction, and hence potentially display different isotopic distributions). At each of these metabolite nodes one contributing reaction involves CO_2_, either as a cleavage or condensation. More specifically, three metabolite nodes involve CO_2_ cleavage (PEP, Pyr, and Suc) and two nodes involve CO_2_ condensation (Mal and OAA). The latter reaction, where a N-1 carbon backbone is converted into a N carbon backbone involving CO_2_ condensation, will be analysed in more detail below.

**Figure 2.**
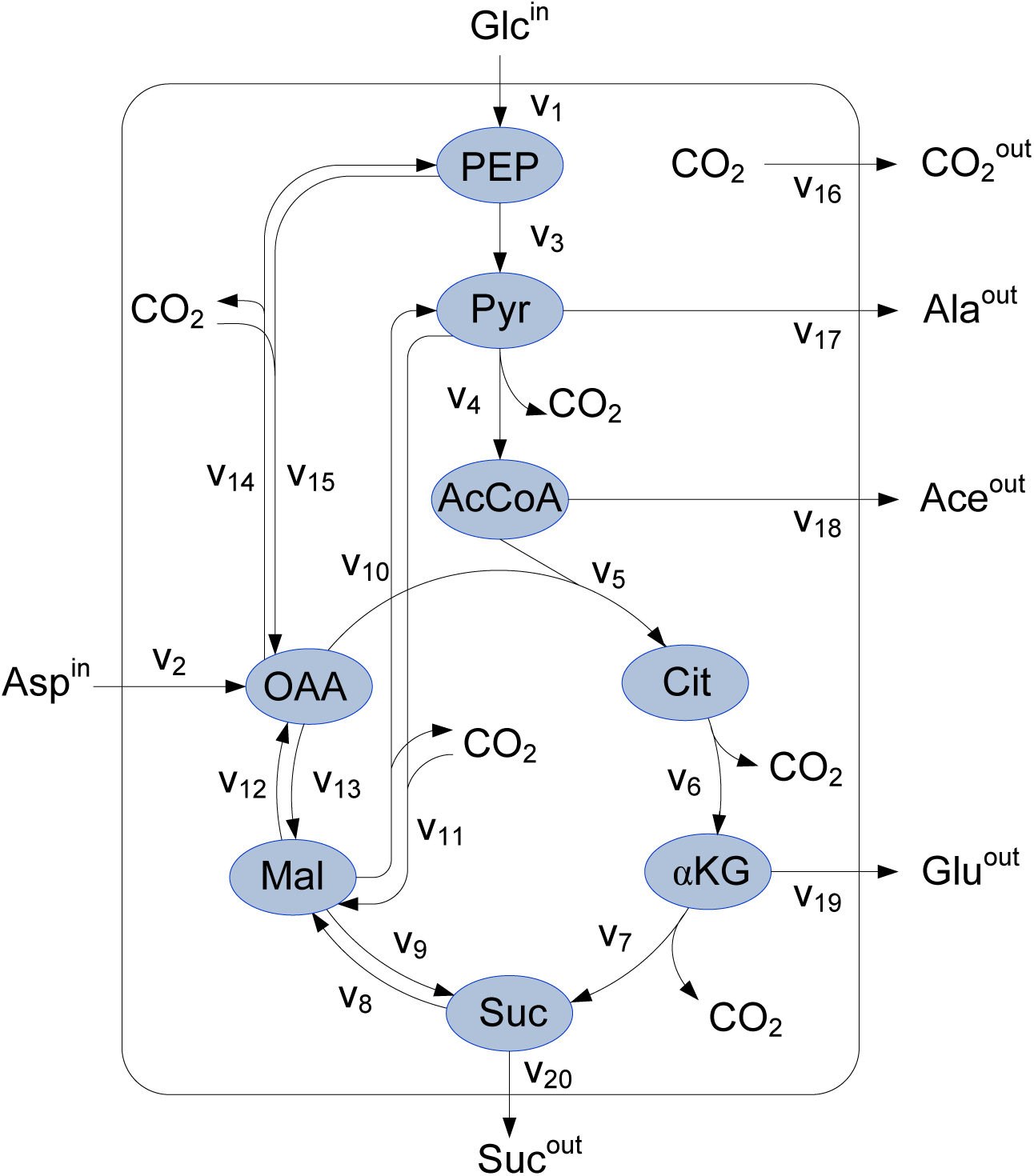
The model reaction network. The network includes the prototypical TCA cycle, lower glycolysis, and several anaplerotic reactions. It is based on central carbon metabolism network topology published by Saunders et al., in 2011 [34]. The solid square represents the system boundary; Glc^in^ and Asp^in^ represent glucose and aspartate, assumed to be only substrates; Suc^out^, Glu^out^, Ace^out^, Ala^out^, and 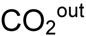 represent secreted succinate, glutamate, acetate, alanine, and carbon dioxide, respectively; PEP, Pyr, AcCoA, Cit, αKG, Suc, Mal and OAA represent intracellular phosphoenolpyruvate, pyruvate, acetyl coenzyme A, citrate, α-ketoglutarate, succinate, malate and oxaloacetate, respectively. The reactions are as given in Table 1.

**Table 1.** The reactions for the model shown in Figures 2 and 3, with their stoichiometries.

| Flux | Description | Stoichiometry |
| --- | --- | --- |
| V<sub>1</sub> | PEP produced from glucose via glycolysis | Glc<sup>in</sup> = PEP |
| V<sub>2</sub> | Aspartate uptake | Asp<sup>in</sup> = OAA |
| V<sub>3</sub> | Pyruvate kinase | PEP = Pyr + ATP |
| V<sub>4</sub> | Pyruvate dehydrogenase complex | Pyr = AcCoA + CO<sub>2</sub> + NADH |
| V<sub>5</sub> | Citrate synthase | AcCoA + OAA = Cit |
| V<sub>6</sub> | Aconitase and Isocitrate dehydrogenase | Cit = αKG + CO<sub>2</sub> + NADPH |
| V<sub>7</sub> | α-ketoglutarate dehydrogenase and succinyl-CoA synthase | αKG = Suc + CO<sub>2</sub> + NADH + ATP |
| V<sub>8</sub> | Succinate dehydrogenase and fumarase | Suc = Mal + CoQ<sub>10</sub>H<sub>2</sub> |
| V<sub>9</sub> | Fumarate hydratase and fumarate reductase | Mal + NADH = Suc |
| V<sub>10</sub> | Malic enzyme | Mal = Pyr + CO<sub>2</sub> + NADPH |
| V<sub>11</sub> | Malic enzyme | Pyr + CO<sub>2</sub> + NADPH = Mal |
| V<sub>12</sub> | Malate dehydrogenase | Mal = OAA + NADH |
| V<sub>13</sub> | Malate dehydrogenase | OAA + NADH = Mal |
| V<sub>14</sub> | PEP carboxykinase | OAA + ATP = PEP + CO<sub>2</sub> |
| V<sub>15</sub> | PEP from glycolysis | PEP + CO<sub>2</sub> = OAA + ATP |
| V<sub>16</sub> | Carbon dioxide secretion | CO<sub>2</sub> = CO<sub>2</sub><sup>out</sup> |
| V<sub>17</sub> | Alanine secretion | Pyr = Ala<sup>out</sup> |
| V<sub>18</sub> | Acetate secretion | AcCoA = Ace<sup>out</sup> |
| V<sub>19</sub> | Glutamate secretion | αKG = Glu<sup>out</sup> |
| V<sub>20</sub> | Succinate secretion | Suc = Suc<sup>out</sup> |

**Figure 3.**
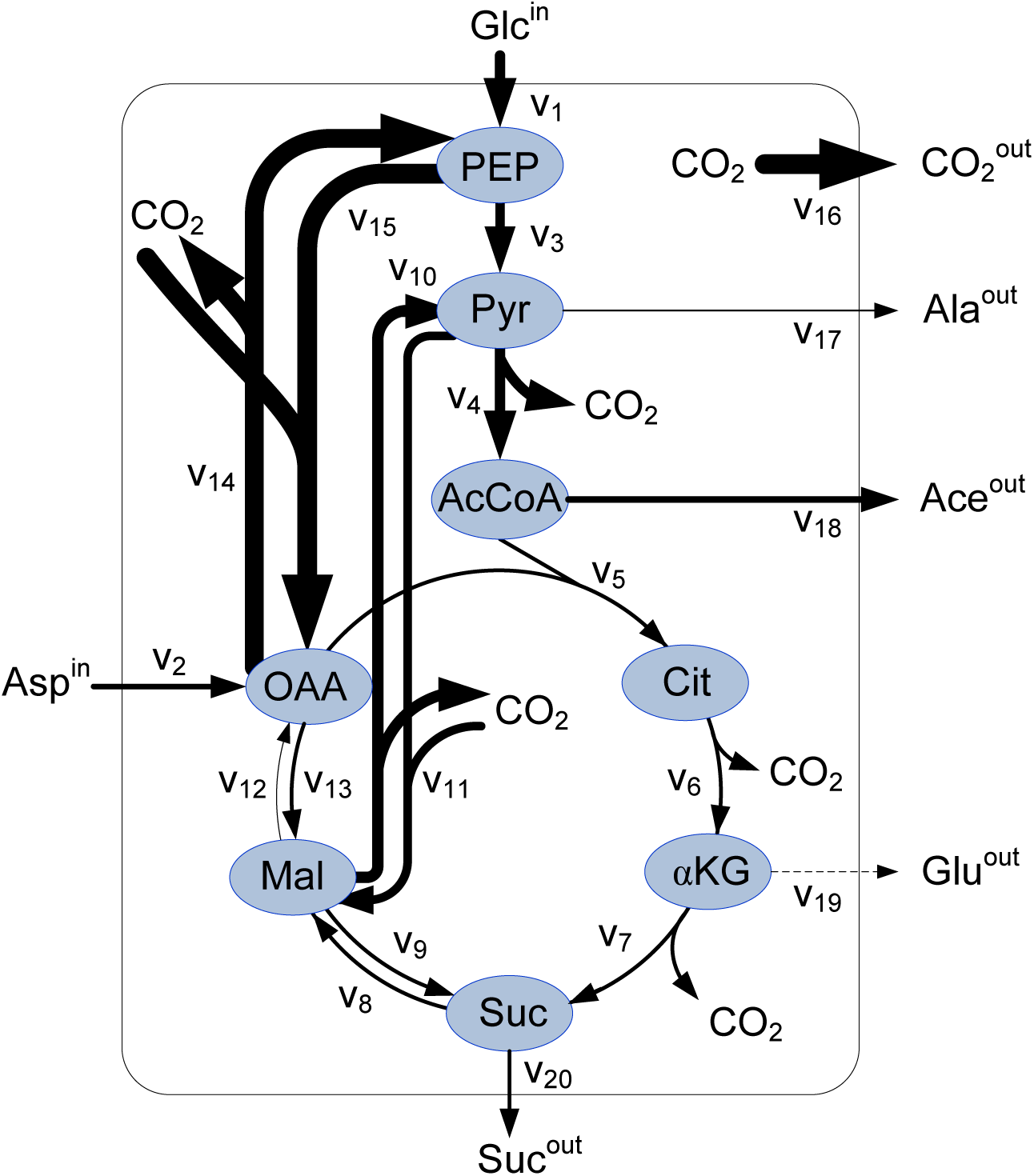
Graphical representation of synthetic fluxes assumed to operate in the reaction network in Figure 2 The assumed fluxes are as follows: v_1_ = 1.1, v_2_ = 0.6, v_3_ = 1.0, v_4_ = 1.1, v_5_ = 0.4, v_6_ = 0.4, v_7_ = 0.4, v_8_ = 0.4, v_9_ = 0.4, v_10_ = 1.2, v_11_ = 0.9, v_12_ = 0.1, v_13_ = 0.4, v_14_ = 2.0, v_15_ = 2.1, v_16_ = 2.1, v_17_ = 0.2, v_18_ = 0.7, v_19_ = 0, v_20_ = 0.4. The zero flux (i.e. v_19_) is shown as a dashed line; the thickness of the lines representing the rest of the fluxes is directly proportional to their value.

Consider a model reaction involving five metabolites D, E, F, G and CO_2_, assumed to be a part of a larger metabolic network, and to combine as shown in Figure 1B. Metabolites D and E are converted to G without a loss or gain or carbon atoms, while the metabolite F combines with CO_2_ to produce G (therefore, metabolites D, E, and G have N carbon atoms, while metabolite F has N-1 carbon atoms). The fractional contributions to the total influx into the node G are:

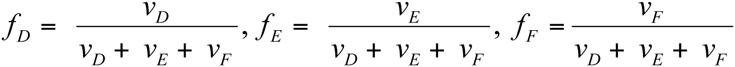

the mass isotopic distribution vector for the metabolite G can be written as:

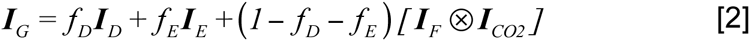

where f_D_ + f_E_ ≤ 1, and **I**_F_ ⊗ **I**_CO2_ represents the discrete convolution of mass isotopic distribution vectors for metabolites F and CO_2_.

Since the elements of mass isotopic distribution vectors are normalized to one, we can replace **I**_CO2_ = [x, 1-x]. If **I**_D_, **I**_E_, and **I**_F_ are measured, the Equation 2 has three unknowns on the right-hand side: f_E_, f_D_, and x. The quantity on the left hand side is the isotopic distribution vector **I**_G_ of the metabolite G. Furthermore, we denote the measured value of **I**_G_ as **I**_Gm_, and define function Φ_1_ such that:

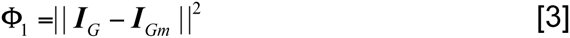

where double bars denote a vector 2-norm. If **I**_D_, **I**_E_, **I**_F_, and **I**_G_ are measured, Φ_1_ is a function of f_E_, f_D_, and x. Hence, the values for f_E_, f_D_, and x can be obtained by constrained optimization of the function Φ_1_ (f_E_, f_D_, x):

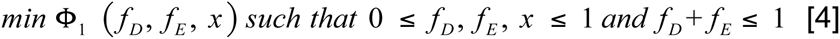

where Φ_1_ is defined by the Equation 3. Typically an optimisation algorithm is employed to solve Equation 4 (e.g. Matlab *FMINCON* function [39]), and the main challenge then is to ensure that the algorithm has reached a global, rather than a local minimum.

### Two metabolite nodes with CO_2_ condensation

The extended version of the previous scenario is depicted in Figure 1C. The relationship between metabolites D, E, F, and G is as described previously. We also have a mirror metabolite node S, which is coupled to the node G via CO_2_. The metabolites P and Q convert to S without a loss or gain or carbon atoms, while the metabolite R combines with CO_2_ to produce G (therefore, metabolites P, Q, and S have N carbon atoms, while metabolite R has N-1 carbon atoms). The mass isotopic distribution vector for the metabolite G is as per Equation 2, and the mass isotopic distribution vector for the metabolite S can be written as:

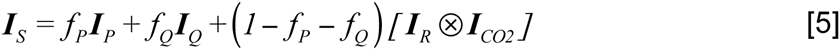

We denote the measured value of **I**_S_ as **I**_Sm_, and define function Φ_2_ such that:

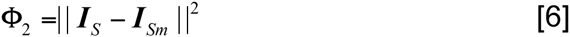

If **I**_P_, **I**_Q_, **I**_R_, and **I**_S_ are measured, Φ is a function of f_p_, f_Q_, and x. And the values for f_P_, f_Q_, and x can be obtained by constrained optimization of the function Φ_2_ (f_A_, f_B_, x):

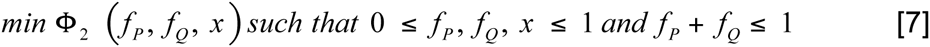

Recognizing that the mass isotopic distribution of CO_2_ is a property of the system that arises from mass balances in the metabolic and isotopic steady-state (Figure 2), one can combine Equations [4] and [7] into a single optimization problem:

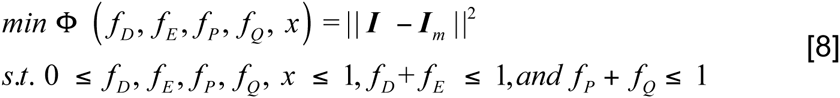

where **I** = [**I**_G_, **I**_S_], **I**_m_ = [**I**_Gm_, **I**_Sm_] (see Equations 2 and 5).

### Parasite strains and labelling conditions

*L. mexicana* wild type (WT) (MNYC/BZ/62/M379) promastigotes were cultured in RPMI medium (Sigma) supplemented with 10% heat inactivated foetal bovine serum (iFBS, GibcoBRL) at 27 °C. Parasites were passaged twice weekly (1/100 and 1/1000) into fresh media to maintain log phase growth. Mid-log phase promastigotes (1 × 10^7^ cell/ml) were harvested two days after inoculation of the media. Cells were pelleted and resuspended in a pre-labelling media to equilibrate (at 2 × 10^7^ cells/mL, 1 hour, 27 °C). Briefly, this completely defined media (modified from [40]) included glucose (6 mM, ^12^C-U-glc), and most other amino acids except for the non-essential amino acid, alanine. Fatty acids were included as lipid containing bovine serum albumin (BSA, 0.5 % final). After equilibration, cells were pelleted (805 × g, 10 minutes) and resuspended in the labelling media containing 3 mM [U-^13^C_6_] glucose and 3 mM [1−^13^C_1_] glucose for 15 hr in order to reach isotopic equilibrium (2 × 10^7^ cells/mL).

### Sampling and metabolite extraction

Parasite metabolism was quenched as described in [41]. Briefly, aliquots of promastigote batch cultures (8 × 10^8^ parasites total, approximately 40 mL) were metabolically quenched by immersing the 75 cm^2^ flask in a dry ice-ethanol slurry to rapidly cool the cell suspension to 0 °C. Immediately, the chilled parasites were centrifuged (805 × g, 10 minutes, 0 °C) and the resulting cell pellet was suspended in ice-cold PBS and 16 replicates (4 × 10^7^ cell total) were transferred to a microfuge tube and washed twice with cold PBS (10,000 × g, 1 min, at 0 °C). The cell pellet was extracted in chloroform:methanol:water (CHCl_3_:CH_3_OH:H_2_O, 1:3:1 v/v, 250 µL, vortex mixed), containing 1 nmol scyllo-inositol as internal standard (60 °C, 15 min). Insoluble material was removed by centrifugation (16,100 × g, 0 °C, 5 minutes) and the supernatant adjusted to CHCl_3_:CH_3_OH:H_2_O 1:3:3 v/v, with addition of H_2_O, vortex mixed and centrifuged (16,000 × g, 5 min) to induce phase separation. The upper phase of the replicates were transferred to a fresh microfuge tube for analysis of polar metabolites.

### Analysis of polar and apolar metabolites by GC-MS

Polar phases were dried into 250 µL glass vial inserts (*in vacuo*, 55 °C) and free aldehyde groups protected by derivitization in methoxyamine chloride (Sigma, 20 µl, 20 mg/ml in pyridine) with continuous mixing (14 hrs, 25 °C). Replicates were derivitized (silylation) with either BSTFA (N,O-bis(trimethylsilyl) trifluoroacetamide containing 1% trimethylchlorosilane (TMCS), Pierce, 20 µl, 1 hr, 25 °C) or MTBSTFA (N-Methyl-N-[tert-butyldimethyl-silyl]trifluoroacetimide containing 1% tert-butyldimethylchlorosilane (TBDMCS), Pierce, 20 µl, 1 hr, 25 °C). All samples were analysed by GC-MS with a DB5 capillary column (J&W Scientific, 30 m, 250 µm i.d., 0.25 µm film thickness), with a 10 m inert duraguard and fitted with a gerstel autosampler. The injector insert and GC-MS transfer line temperatures were 270 °C and 250 °C, respectively. The oven temperature gradient was programmed as follows: 70 °C (1 min); 70 °C to 295 °C (12.5 °C/min); 295 °C to 320 °C (25 °C/min); 320 °C (2 min). Data was collected in both SCAN and selected ion-monitoring (SIM) modes for metabolite identification and quantification (dwell time 20 ms), respectively. For TMS derivitised samples, the metabolites and ions monitored were: PEP (369.1, 370.1, 371.1, 372.1, and 373.1 *m/z*), citrate (465.15, 466.15, 467.15, 468.15, 469.15, 470.15, 471.15, and 472.15 *m/z*), succinate (247.1, 248.1, 249.1, 250.1, 251.1 and 252.1 *m/z*), and malate (335.1, 336.1, 337.1, 338.1, 339.1, and 340.1 *m/z*). For TBDMS derivitised samples, the metabolites and ions monitored were: alanine (260, 261, 262, and 263 *m/z*), aspartate (418, 419, 420, 421, and 422 *m/z*) and glutamate (432, 433, 434, 435, and 436 *m/z*). For the TBDMS samples, SIM data was collected in split/splitless mode Replicates were randomised and similarly prepared standards were also analysed.

## Results

### Validation using a model network

A simple model network (shown in Figure 2), comprising of lower glycolysis (v_1_ and v_3_), TCA cycle (v_4_-v_8_, v_12_), glycosomal succinate fermentation (v_9_, v_13_, and v_15_) pathway [42] and several anaplerotic reactions (v_10_, v_11_, v_14_), was constructed based on previously published data for *L. mexicana* [34,37]. Glucose and aspartate are assumed to be the only carbon sources (alanine was not present in the media). And carbon leaves the network via carbon dioxide, alanine, acetate, glutamate, and succinate. There are a total of 20 fluxes in this reaction network, and assuming a metabolic steady-state it results in nine equations as shown in Table 2. Assuming flux values for the reactions participating in the network and isotopic composition of input substrates it is possible to calculate exact mass isotopic distributions of individual metabolites at metabolic and isotopic steady-state. To validate the method for calculating flux ratios described above, we assumed the reaction fluxes shown in Figure 3, the composition of input substrates, and calculated the steady-state mass isotopic distributions for the carbon backbone of phosphoenolpyruvte (PEP), pyruvate (Pyr), succinate (Suc), malate (Mal), oxaloacetate (OAA), and CO_2_ in metabolic and isotopic steady-state (see Table 3). Then we applied the Equation 4 to back-calculate the fractional fluxes around Mal and OAA nodes.

**Table 2.** Metabolic steady-state equations for the network shown in Figures 2 and 3.

| Node | Metabolic steady-state equation |
| --- | --- |
| PEP | V<sub>1</sub> + V<sub>14</sub> = V<sub>3</sub> + V<sub>15</sub> |
| Pyr | V<sub>3</sub> + V<sub>10</sub> = V<sub>4</sub> + V<sub>11</sub> + V<sub>17</sub> |
| AcCoA | V<sub>4</sub> = V<sub>5</sub> + V<sub>18</sub> |
| Cit | V<sub>5</sub> = V<sub>6</sub> |
| αKG | V<sub>6</sub> = V<sub>7</sub> + V<sub>19</sub> |
| Suc | $V_7 + V_9 = V_8 + V_{20}$ |
| Mal | $V_8 + V_{11} + V_{13} = V_9 + V_{10} + V_{12}$ |
| OAA | $V_2 + V_{12} + V_{15} = V_5 + V_{13} + V_{14}$ |
| CO<sub>2</sub> | $V_4 + V_6 + V_7 + V_{10} + V_{14} = V_{11} + V_{15} + V_{16}$ |

**Table 3.** Calculated steady-state mass isotopic distributions for the carbon backbone of the key metabolites for the assumed fluxes in Figure 3.

|  | m<sub>0</sub> | m<sub>1</sub> | m<sub>2</sub> | m<sub>3</sub> | m<sub>4</sub> |
| --- | --- | --- | --- | --- | --- |
| CO<sub>2</sub> | 0.747152 | 0.252848 |  |  |  |
| PEP<sub>(3)</sub> | 0.447893 | 0.189819 | 0.017475 | 0.344812 |  |
| Pyr<sub>(3)</sub> | 0.453369 | 0.200297 | 0.052123 | 0.294211 |  |
| Asp<sub>(4)</sub><sup>in</sup> | 0.957882 | 0.041441 | 0.000672 | 0.000005 | 0.000000 |
| Suc<sub>(4)</sub> | 0.314743 | 0.220152 | 0.226117 | 0.159334 | 0.079653 |
| Mal<sub>(4)</sub> | 0.363796 | 0.240843 | 0.112386 | 0.208842 | 0.074134 |
| OAA<sub>(4)</sub> | 0.469236 | 0.208786 | 0.049947 | 0.203994 | 0.068036 |

It is straightforward to show that for the assumed fluxes in Figure 3, the steady-state equations hold (Table 2). The substrate labelling was assumed to be 50% [U-^13^C_6_] + 50% [1−^13^C_1_] glucose and unlabelled aspartate (Glc^in^ and Asp^in^ in Figure 3). Mass spectrometry measurements yield mass isotopic distributions of molecular fragments that can be corrected to obtain mass isotopic distribution vectors of the carbon backbone alone. Therefore for the purpose of this work, the values in Table 3 represent idealized measurements (no experimental error), obtained on the metabolic network shown in Figure 2 and a specific set of fluxes (Figure 3).

For Mal node, Equations 3 and 4 can be written explicitly as follows:

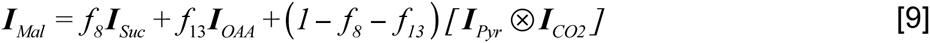

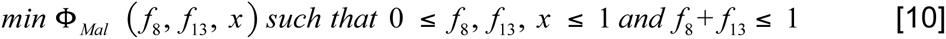

where f_8,_ f_11_, and f_13_ are fractional fluxes corresponding to net fluxes v_8_, v_11_, and v_13_ respectively (see Figure 2), and f_11_ = 1 – f_8_ – f_13_. Similarly, for OAA node Equations 3 and 4 can be written as:

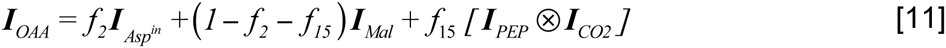

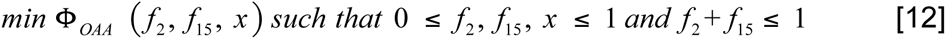

where f_2_, f_12_, and f_15_ are fractional fluxes corresponding to v_2_, v_12_, and v_15_, respectively (Figure 2), and f_12_ = 1 – f_2_ – f_15_. In the above equations, the variable x refers to m_0_ in the mass isotopic distribution vector of CO_2_ (i.e. **I**_CO2_ = [x, 1-x], see above). As before, recognizing that the mass isotopic distribution of CO_2_ is a property of the system that arises from mass balances in the metabolic and isotopic steady-state, one can combine Equations [10] and [12] into a single optimization problem:

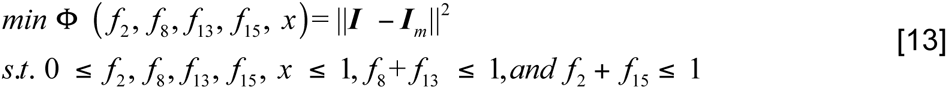

where I = [I_Mal_, I_OAA_] is the concatenation of calculated mass isotopic distribution vectors for Mal and OAA respectively, Equations 9 and 11 or explicitly:

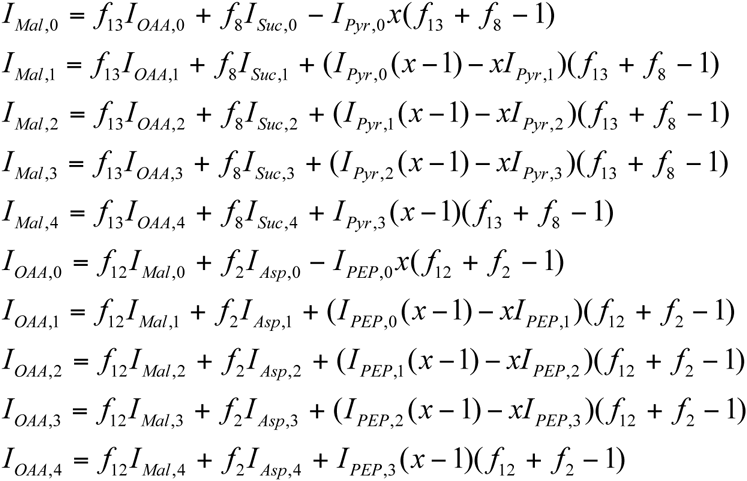

and I_m_ is the concatenation of the same measured vectors.

This formulation of the problem allows a single optimization of the objective function of five variables, subject to upper and lower boundary constraints as well as two independent inequality constraints, to obtain fractional distribution of incoming fluxes around Mal and OAA nodes. Since the proposed approach delineates flux distributions around two nodes simultaneously, it falls between the traditional ^13^C MFA (resolves all fluxes in the underlying network topology) and metabolic flux ratio analysis (resolves distribution of fluxes around a single node).

### Stability of the solution

We investigated stability of the optimization described by Equation 13, by repeated optimizations on idealized data. The fluxes were assumed as shown in Figure 3, and an equimolar mix of [U-^13^C_6_] and [1−^13^C_1_] glucose and unlabelled aspartate were assumed as substrates. Isotopic steady-state labelling of PEP, Pyr, Suc, Mal, and OAA were calculated by EMU methodology [43]. The calculated values of mass isotopic vectors were confirmed by forward labelling calculations carried in OpenFlux [44] and Metran [45]. We employed 100 randomly initialised runs of Matlab *FMINCON* optimiser to compute the fractional fluxes for the malate and oxaloacetate nodes. The results (Figure 4) showed that the sum of squared residuals (SSR) value returned was different each time. This means that either objective function surface is very flat causing the optimisation algorithm to terminate in a vicinity of the global minimum (but rarely, if ever, at the global minimum) or the surface of the objective function had numerous closely spaced shallow local minima in the vicinity of the global minimum. Moreover, the existence of the global minimum is not guaranteed. Further tests with 1000, 10000, 100000 and 1000000 randomly initialised runs of *FMINCON* optimiser were conducted, and although each time the results got sufficiently close to the expected (synthetic) values, SSR value distribution was similar to that of 100 runs (Figure 4). In the case of experimental data (where the results are not known) it would have been very difficult to ascertain how many randomly initialised optimisation runs were sufficient. Decreasing the value of the objective function stopping criteria and increasing the number of iterations per optimisation run was also tested with unsatisfactory results. Noting that if x was known both fractional fluxes could be solved for analytically via Equation 1, an alternative approach is to vary x from 0 to 1, compute fractional fluxes and the objective function Φ, and then plot the value of Φ against the value of x. This will reveal the existence and the location of the global minimum, and the corresponding values. Firstly, we utilised quadratic programming to calculate the objective function value for a range of, for example, 100,000 equidistant x values. The results are shown in Figure 5A, and they indicate that the objective function has a global minimum. The minimum value of this new range is shown in Figure 5B, and has been observed to produce more precise results than 1, 000, 000 runs of *FMINCON* optimiser in the previous section (Table 4). Moreover, on a computer where 1, 000, 000 runs of *FMINCON* optimiser takes 10 hours, this approach runs in less than 10 minutes and is therefore suitable for high-throughput applications.

**Figure 4.**
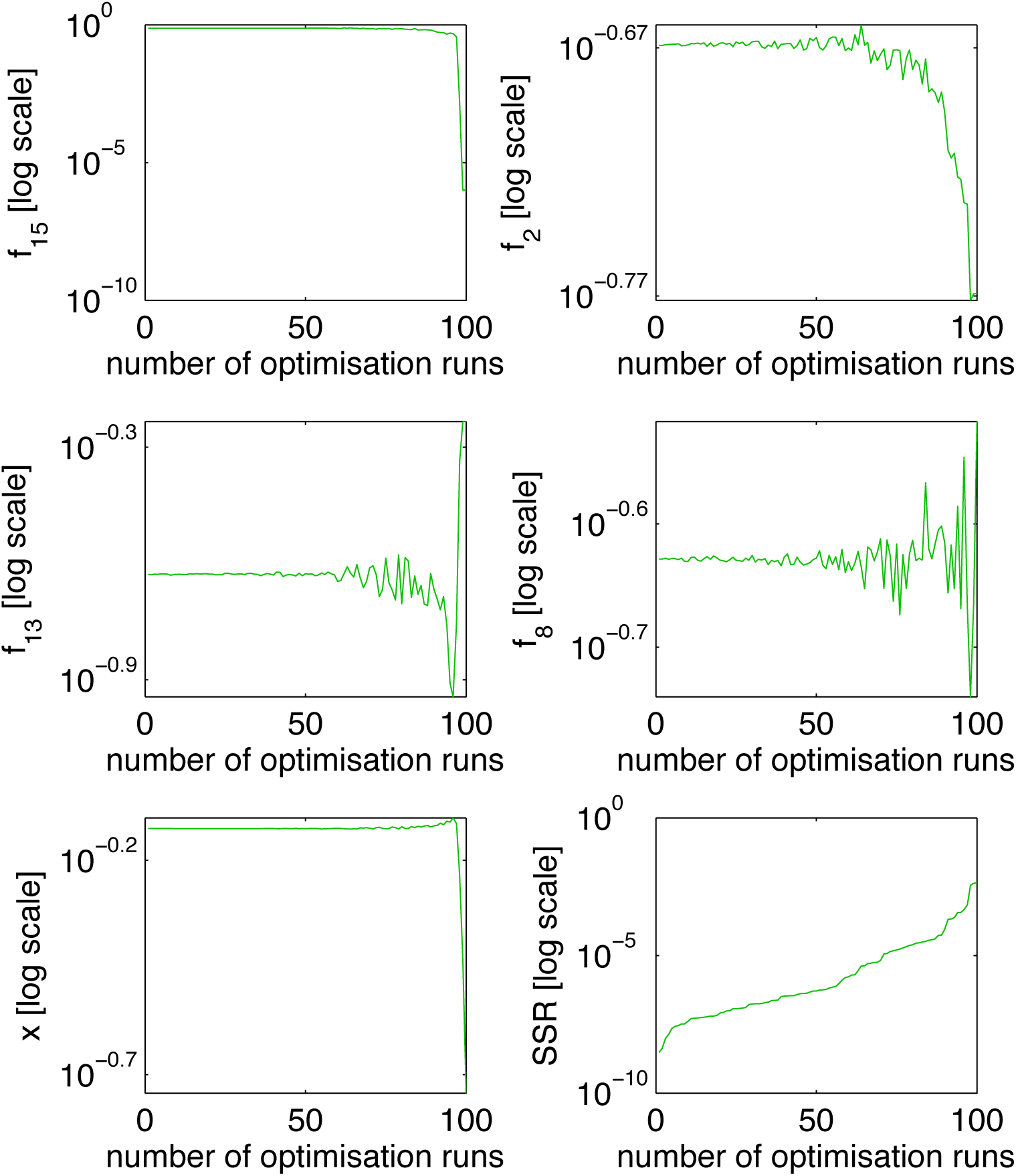
Results based on ideal synthetic data and 100 runs of Matlab’s Optimisation run numbers are based on the results sorted by the value of the objective function (i.e. 1^st^ optimisation run has the smallest objective function value, and 100^th^ optimisation run has the largest objective function value. Similar results were obtained for 1000, 10000, 100000 and 1000000 optimisation runs, in that no single value for the objective function is reached in more than one run.

**Figure 5.**
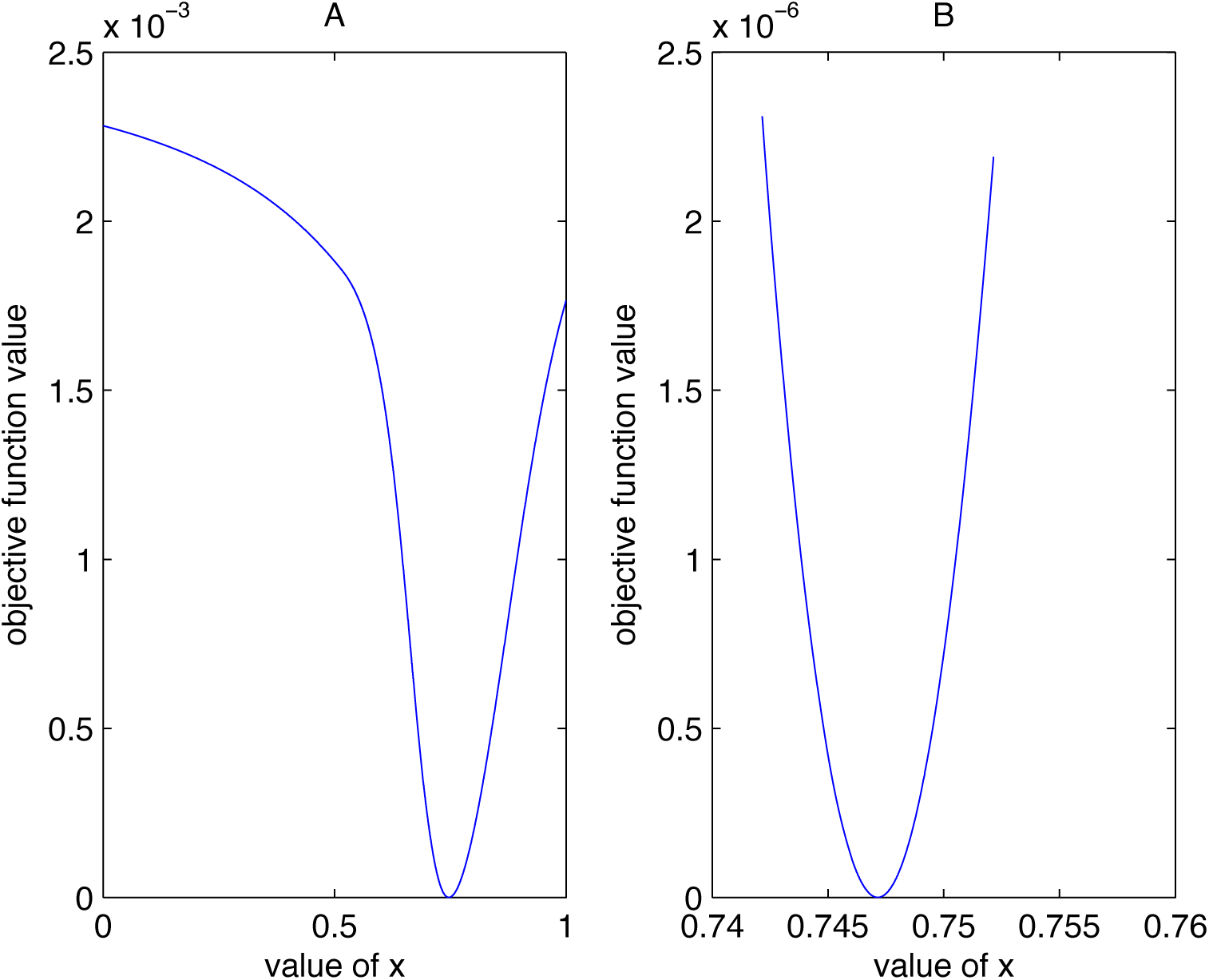
Results based on ideal synthetic data without an optimisation function. Panel (A) shows the value of the objective function for 1000 equidistant values of x. Panel (B) zooms in on 10 of the previous points belonging to the values of x closest to the x value corresponding to the objective function minimum.

**Table 4.** Comparison of solutions with and without the use of the optimisation function.

| Variable | Actual values | *FMINCON* | *QUADPROG* |
| --- | --- | --- | --- |
| SSR | 0.0000 | $4.89 \cdot 10^{-11}$ | $2.05 \cdot 10^{-12}$ |
| f<sub>8</sub> | 0.2353 | 0.235320 | 0.235298 |
| f<sub>13</sub> | 0.2353 | 0.235298 | 0.235281 |
| f<sub>2</sub> | 0.2143 | 0.214273 | 0.214281 |
| f<sub>15</sub> | 0.7500 | 0.749971 | 0.749979 |

Comparing our approach (without the use of an optimisation function) with Matlab's *FMINCON* optimisation function. The second column shows the best of 1,000,000 randomly initialised *FMINCON* optimisation runs. The third column shows the values obtained via the quadratic programming approach. Note that the values in both columns were produced using ideal synthetic data.

### Validation in the presence of noise

Experimental data will always be affected by noise, various artifacts due to machine imperfections, and possibly human error. Therefore an important question is, are the fractional fluxes computable in the presence of realistic levels of experimental noise? In instruments that deploy detectors with electron multipliers, a proportional relationship between the measured mean intensity and variance is often observed (e.g. in mass spectrometry [46] and microarrays [47]. In our data (five metabolites, a total of 23 *m/z* channels measured), a strong relationship between the mean intensity and variance was not apparent (Figure 6). The observed correlation coefficient between the mean intensity and variance was 0.448, and the p-value for testing the hypothesis of no correlation was 0.03. Furthermore, the data for each *m/z* channel appeared normally distributed in the first approximation: for only one out of 23 *m/z* channels the hypothesis that the observed values are drawn from a normal distribution could be rejected at the 95% confidence level by Lilliefors test (specifically for Mal m_0_+2 data set, which showed two groups of tightly grouped values). However, the distribution of variances calculated for 23 *m/z* channels (Figure 6, y-axis) failed Kolmogorov-Smirnov and Lilliefors tests for normality. Therefore, the observed distribution of variances across different *m/z* channels is unlikely to be normal. To approximate this distribution, we fitted the parameters of the Weibull distribution to the calculated variances across 23 *m/z* channels. The Weibull distribution is widely used in reliability engineering and life data analysis because it can model a wide range of behaviours [48]. Best fit to the experimental data resulted in α=0.0031 (scale) and β=1.5315 (shape) of the Weibull distribution. Therefore to simulate noise similar to that observed experimentally, we assumed the following: (a) no correlation between the noise variance and m/z channel intensity; (b) noise for each *m/z* channel was normally distributed; and (c) the variances of individual *m/z* channels follow the Weibull distribution with α=0.0031 and β=1.5315.

**Figure 6.**
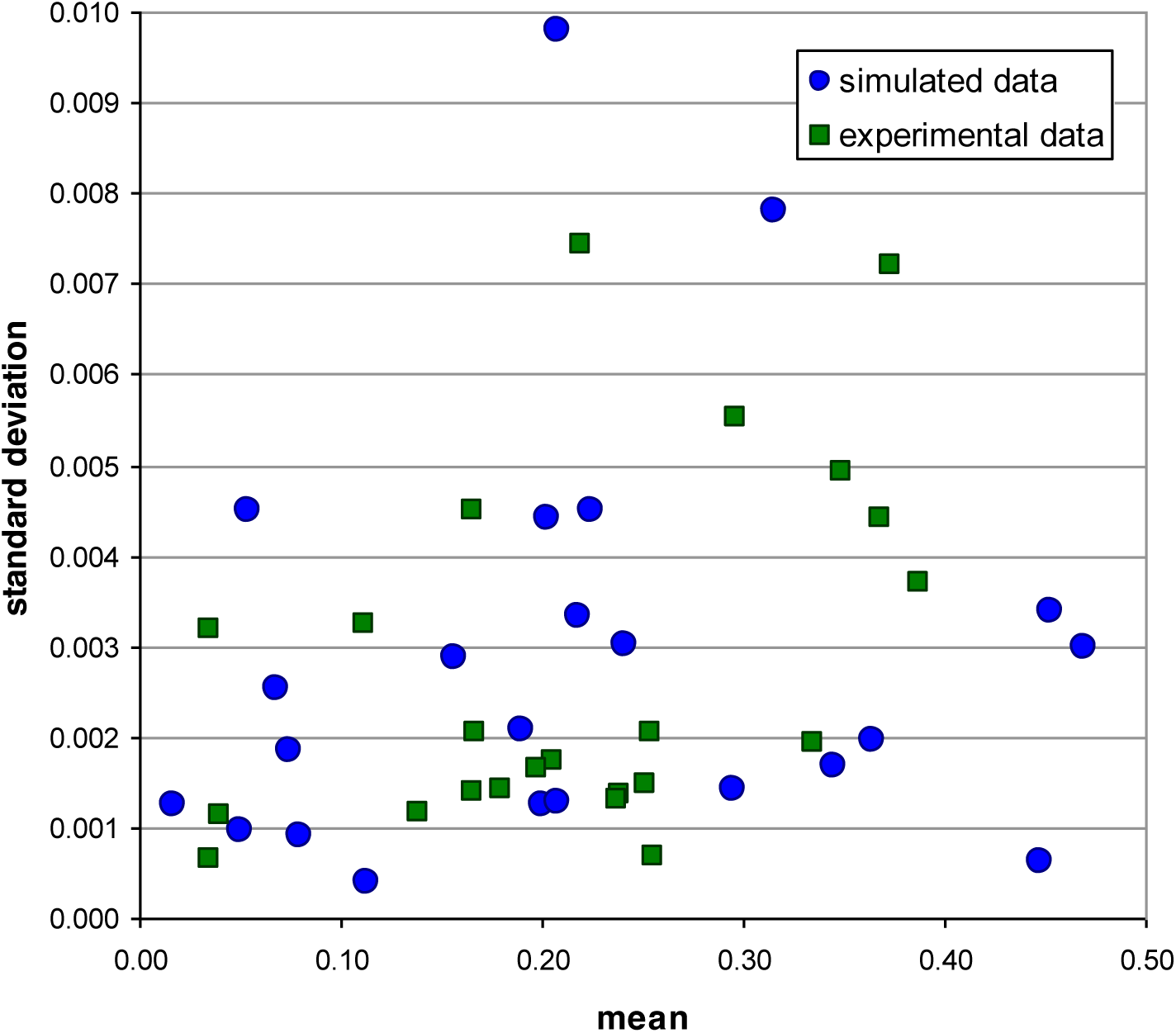
Intensity mean vs. standard deviation for 23 m/z channels across five metabolites. Intensity mean vs. standard deviation for 23 m/z channels across five metabolites (PEP, Pyr, Mal, Suc, and OAA). For experimental data plotted are the mean and standard deviation calculated from eight replicate experiments. For the simulated data set, *m/z* channel intensities were taken from mass isotopic distributions of five metabolites calculated based on fluxes shown in Figure 3, and standard deviations were derived from Weibull distribution whose parameters were derived from the best fit to experimental data. Only a weak correlation between the mean intensity and variance in the experimental data was observed (a correlation coefficient of 0.448). The plot suggests that the strategy adopted for the simulation of noise in data approximates well noise in experimental data.

In summary, the data with noise was simulated in the following way. Isotopic steady-state labelling of PEP, Pyr, Suc, Mal, and OAA were calculated by EMU methodology [43], assuming the fluxes as given in Figure 3, and an equimolar mix of [U-^13^C_6_] and [1-^13^C_1_] glucose and unlabelled aspartate as substrates. This produced backbone labelling patterns of PEP, Pyr, Suc, Mal, and OAA, as idealized intensities for 23 *m/z* channels (m_0_, m_0_+1, m_0_+2, m_0_+3 for three carbon backbones of Pyr and PEP, and m_0_, m_0_+1, m_0_+2, m_0_+3, m_0_+4 for four carbon backbones of Suc, Mal, and OAA). To simulate noise, a set of variances was drawn from the Weibull distribution with α=0.0031 and β=1.5315, and variances were randomly assigned to 23 *m/z* channels. In the next step, for each *m/z* channel a noise value was drawn from a normal distribution characterized with the assigned variance, and the noise was added to the channel intensity. The plot of mean intensity vs standard deviation for the experimental data, as well as for an example simulated data with noise, is shown in Figure 6. The fractional flux estimation method (Equation 13) was run with the noisy synthetic data, and it was able to retrieve the original fractional fluxes to within the confidence intervals (Table 5). This validates the approach in the presence of the level of noise found in our experimental data.

**Table 5.** Synthetic fractional flux estimation in the presence of noise.

|  | Oxaloacetate node |  |  | Malate node |  |  |  |
| --- | --- | --- | --- | --- | --- | --- | --- |
|  | f<sub>12</sub> | f<sub>2</sub> | f<sub>15</sub> | f<sub>13</sub> | f<sub>8</sub> | f<sub>11</sub> | SSR |
| actual | **0.0357** | **0.2143** | **0.7500** | **0.2353** | **0.2353** | **0.5294** |  |
| #1 | 0.0495 | 0.2112 | 0.7393 | 0.2185 | 0.2352 | 0.5462 | $3.21 \cdot 10^{-05}$ |
| #2 | 0.1031 | 0.2130 | 0.6840 | 0.2226 | 0.1819 | 0.5955 | $1.14 \cdot 10^{-04}$ |
| #3 | 0.0087 | 0.2140 | 0.7773 | 0.1957 | 0.1820 | 0.6223 | $2.20 \cdot 10^{-05}$ |
| #4 | 0.0000 | 0.2204 | 0.7796 | 0.2490 | 0.2568 | 0.4943 | $9.55 \cdot 10^{-05}$ |
| #5 | 0.0026 | 0.2263 | 0.7712 | 0.2873 | 0.2707 | 0.4420 | $2.36 \cdot 10^{-05}$ |
| #6 | 0.0746 | 0.2135 | 0.7119 | 0.2202 | 0.2473 | 0.5326 | $1.06 \cdot 10^{-05}$ |
| #7 | 0.0516 | 0.2206 | 0.7277 | 0.2465 | 0.2386 | 0.5149 | $2.15 \cdot 10^{-05}$ |
| #8 | 0.0449 | 0.2070 | 0.7481 | 0.2545 | 0.2559 | 0.4896 | $8.84 \cdot 10^{-06}$ |
| **mean** | **0.0419** | **0.2157** | **0.7424** | **0.2368** | **0.2336** | **0.5297** |  |
| **stdev** | **0.0366** | **0.0062** | **0.0339** | **0.0283** | **0.0337** | **0.0585** |  |

The results shown are based on simulated noisy data (8 synthetic replicates).

### Application to experimental data

We applied the optimizaition via Equation 13 to determine fractional fluxes around Mal and OAA nodes in *L. mexicana*. Rapidly dividing *L. mexicana* parasites were cultured in completely defined media containing equimolar [U-^13^C_6_] and [1-^13^C_1_] glucose for 15 hrs. Metabolites were extracted, derivitised and analysed by GC-MS. The normalized data obtained for PEP, Pyr, Suc, Mal, and OAA, corrected for the presence of natural isotopes and heteroatoms [49] is shown in Table 6. For these five metabolites a total of 23 *m/z* channels were measured (m_0_, m_0_+1, m_0_+2, m_0_+3 for three carbon backbones of Pyr and PEP; and m_0_, m_0_+1, m_0_+2, m_0_+3, m_0_+4 for four carbon backbones of Suc, Mal, and OAA). Pyr labelling was measured via free intracellular alanine, and OAA labelling via intracellular aspartate. A total of 8 experimental replicates were measured. The experimental data acquired on *L. mexicana* is shown in Table 6. We assumed the network topology shown in Figure 2, and proceeded to determine fractional fluxes around Mal and OAA nodes according to Equation 13.Table 7 lists best solutions for f_2_, f_8_, f_13_, and f_15_, for each experimental replicate achieved by minimization of the objective function. A simple measure of the uncertainty in the solution is given as standard deviation in Table 7, calculated from eight experimental replicates. The estimated fractional fluxes for Mal node (f_8_, f_11_, and f_13_) and OAA node (f_2_, f_12_, and f_15_) are shown graphically in Figure 7, and a pictorial representation of the fractional fluxes in *L. mexicana* determined from the experimental data is shown in Figure 8. We observed that OAA is chiefly derived from malate (f_12_, 86%) rather then from aspartate (f_2_, 4%) or PEP (f_15_, 10%). Malate was primarily derived from succinate (f8, 61%) and OAA (f_13_, 38%) while pyruvate contribution was found to be negligible (f_11_, 1%).

**Figure 7.**
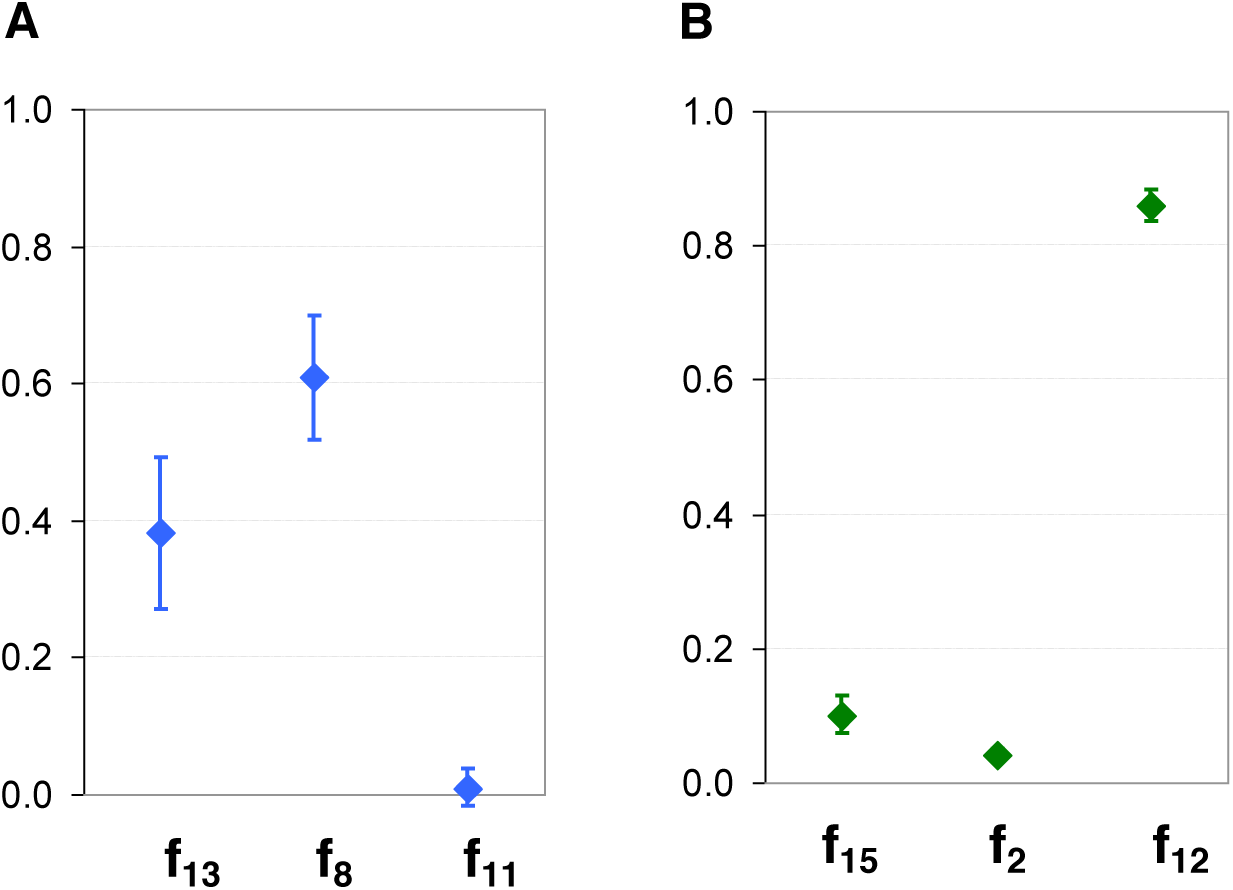
Fractional fluxes f_13_, f_8_, f_11_ (Mal node) and f_15_, f_2_, f_12_ (OAA node). Fractional fluxes f_13_, f_8_, f_11_ (Mal node) and f_15_, f_2_, f_12_ (OAA node) calculated by optimization via Equation 11 for rapidly dividing *L. mexicana* promastigotes. Mean and standard deviation from eight replicate experiments are shown.

**Figure 8.**
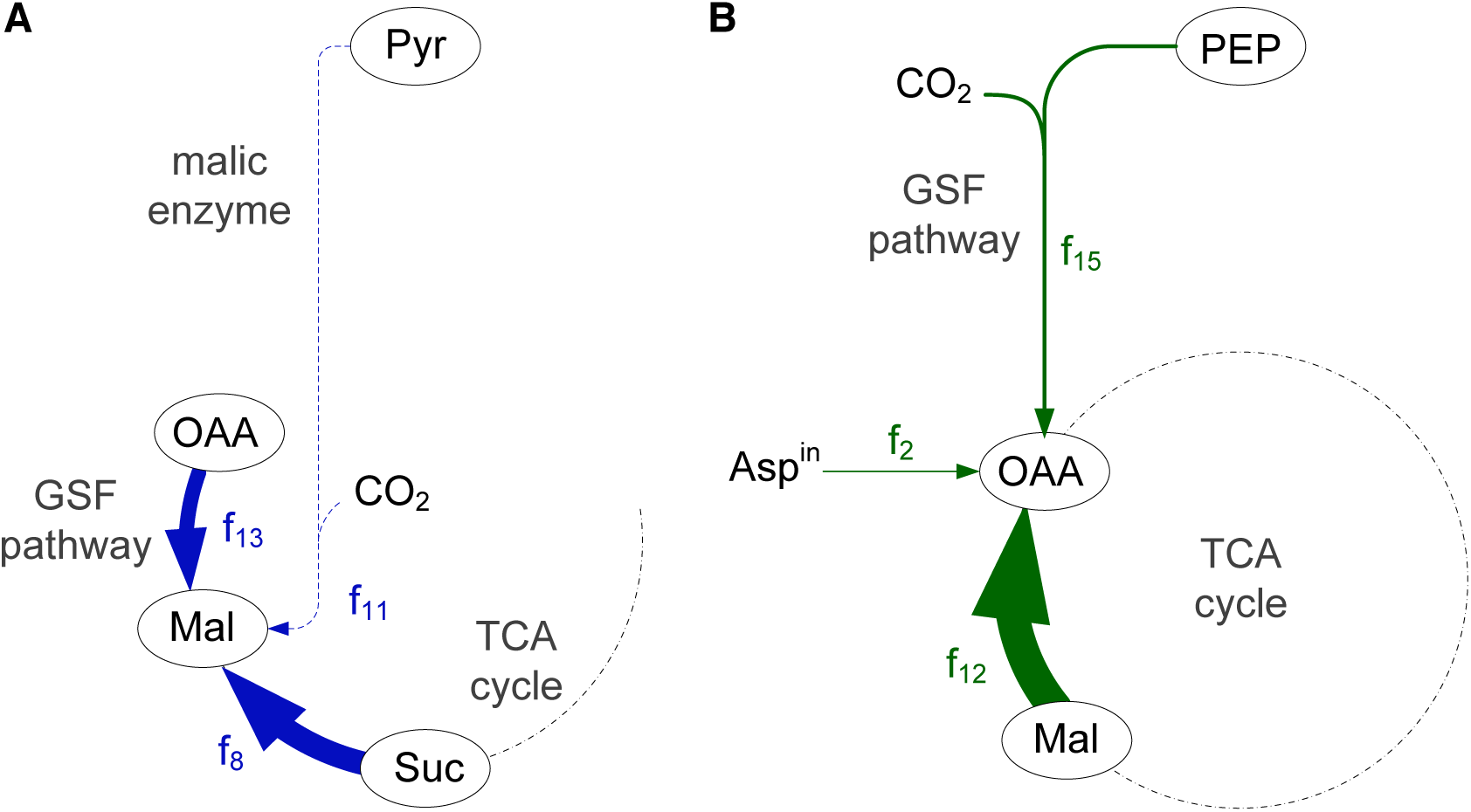
Incoming fractional fluxes around OAA and Mal nodes obtained from experimental data. Shown is the mean based on eight independent replicate experiments. The thickness of the reaction arrows is proportional to the obtained mean values of fractional fluxes inTable 6. Graphical representation is derived from Figure 2 by focusing only on the relevant metabolic subnetwork, all other metabolites and reactions are omitted.

**Table 6.**
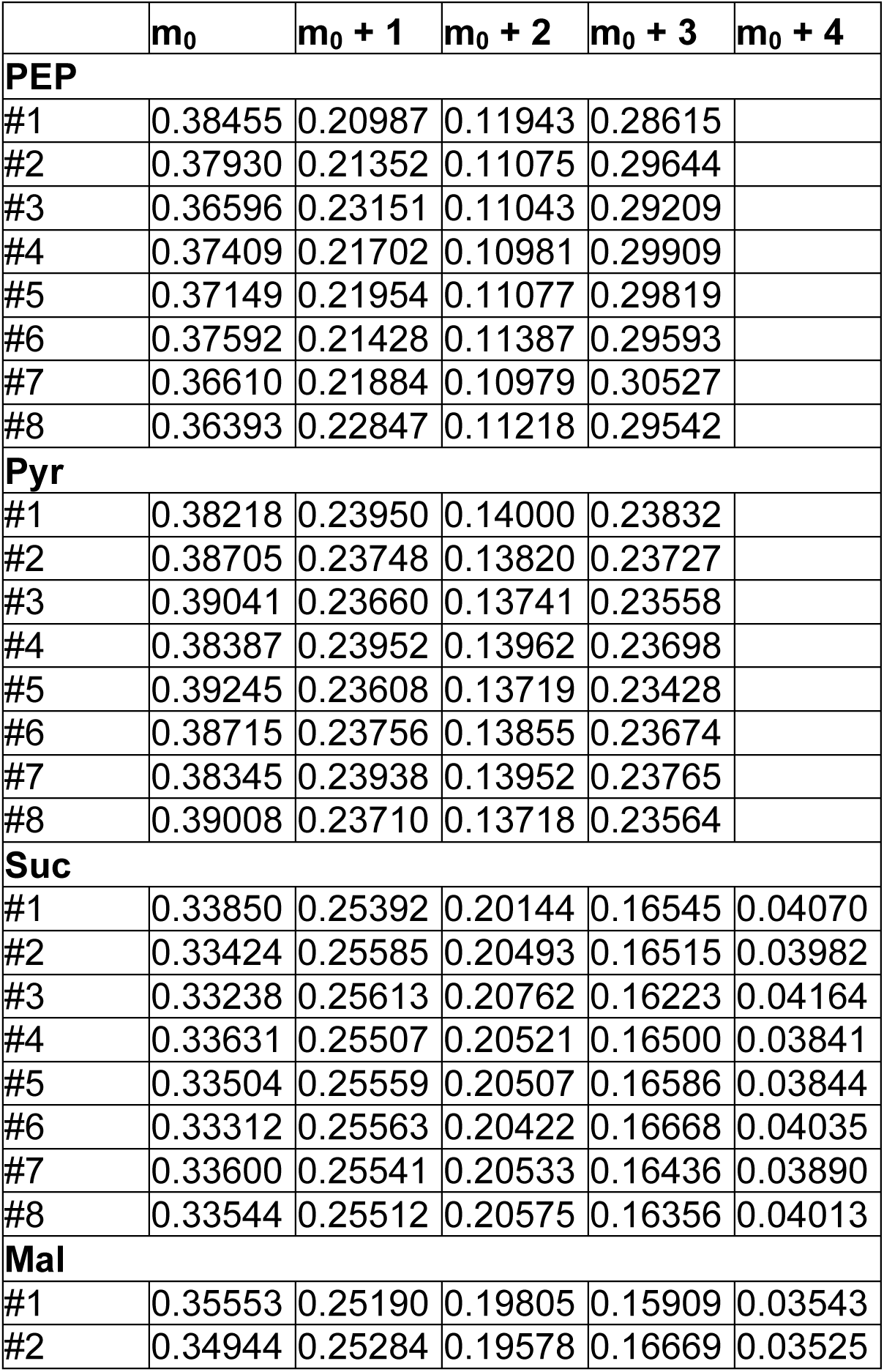
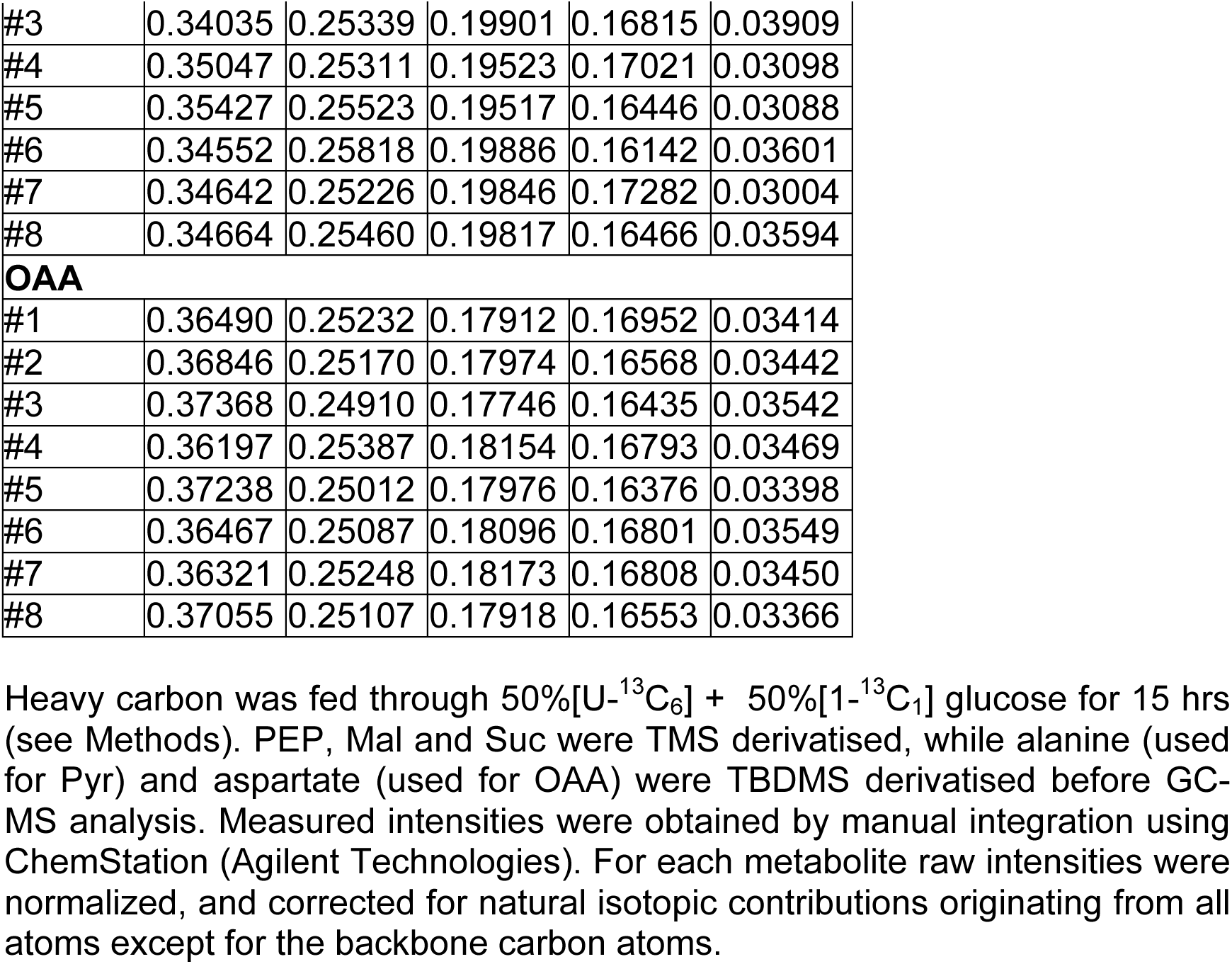
Mass isotopic distributions for PEP, Pyr, Suc, Mal, and OAA in rapidly dividing *Leishmania mexicana* promastigotes at isotopic and metabolic steady state measured by GC-MS.

**Table 7.** Flux ratio estimation results in rapidly dividing *L. mexicana*

|  | **Oxaloacetate node** |  |  | **Malate node** |  |  |  |
| --- | --- | --- | --- | --- | --- | --- | --- |
|  | **f<sub>12</sub>** | **f<sub>2</sub>** | **f<sub>15</sub>** | **f<sub>13</sub>** | **f<sub>8</sub>** | **f<sub>11</sub>** | **SSR** |
| #1 | 0.8172 | 0.0246 | 0.1582 | 0.4271 | 0.5729 | 0.0000 | 1.13·10<sup>-04</sup> |
| #2 | 0.8827 | 0.0384 | 0.0789 | 0.4257 | 0.5743 | 0.0000 | 4.05·10<sup>-05</sup> |
| #3 | 0.8611 | 0.0560 | 0.0829 | 0.1694 | 0.7523 | 0.0783 | 4.84·10<sup>-05</sup> |
| #4 | 0.8846 | 0.0339 | 0.0816 | 0.5190 | 0.4810 | 0.0000 | 5.26·10<sup>-05</sup> |
| #5 | 0.8630 | 0.0440 | 0.0930 | 0.4821 | 0.5179 | 0.0000 | 2.92·10<sup>-05</sup> |
| #6 | 0.8353 | 0.0377 | 0.1270 | 0.3268 | 0.6732 | 0.0000 | 3.98·10<sup>-05</sup> |
| #7 | 0.8701 | 0.0445 | 0.0854 | 0.3925 | 0.6075 | 0.0000 | 9.43·10<sup>-05</sup> |
| #8 | 0.8568 | 0.0450 | 0.0983 | 0.3129 | 0.6871 | 0.0000 | 3.33·10<sup>-05</sup> |
| **mean** | **0.8589** | **0.0405** | **0.1006** | **0.3819** | **0.6083** | **0.0098** |  |
| **stdev** | **0.0229** | **0.0093** | **0.0279** | **0.1108** | **0.0910** | **0.0277** |  |

## Discussion

A total of six flux ratios into the oxaloacetate and malate pools, as well as the mass isotopic distribution of carbon dioxide, have been computed in the rapidly dividing *L. mexicana* promastigotes at metabolic and isotopic steady state (Table 7). These results, shown in Figures 7 and 8, represent the first *in vivo* quantitative fractional flux values for *Leishmania spp.* or any other protozoan. They are also, to the best of our knowledge, the first fractional flux results based on the mass isotopic distribution measurements of free intracellular metabolic intermediates. Together these two firsts highlight the yet unexplored potential of combining ^13^C-MFRA and direct metabolomics measurements for the quantitative study of metabolism. The newly developed approach for simultaneous calculation of *in vivo* fractional fluxes into two or more nodes with carbon dioxide condensation falls in between traditional ^13^C-MFA (computes a number of net fluxes in the network) and ^13^C-MFRA (computes fractional fluxes around a single node). The metabolic reactions where carbon dioxide from the environment is incorporated into a metabolite’s carbon skeleton are commonly encountered in biology. Specifically, the new approach will be essential in experimental set-ups where the carbon dioxide mass isotopic distribution cannot be accurately and precisely quantified (e.g. bioreactors operated in batch mode).

The fractional flux results show that the oxaloacetate pool is predominantly topped up from the malate pool (∼86% of the oxaloacetate carbon backbone comes from the malate carbon skeleton), a small amount (∼10%) of the oxaloacetate is generated from PEP (produced by glycolysis) through the glycosomal succinate fermentation (GSF) pathway, and a very small amount of the oxaloacetate (∼4%) comes from the uptake of exogenous aspartate (Figure 8).

Fractional flux results for the malate node indicate that the malic enzyme operation in the direction from pyruvate to malate is negligible, and almost twice as much malate is created through the TCA cycle from succinate than via the GSF pathway from oxaloacetate. This result confirms the irreversibility of the malic enzyme reaction reported previously [34]. Fractional influxes into the malate pool display considerably larger confidence intervals than fractional influxes into the oxaloacetate pool (Figure 7). Analysis of additional metabolites, such as the mass isotopic distribution of fumarate which is closer to malate than the measured succinate, may improve the fit for the malate node. Alternatively, the larger confidence may actually arise from metabolite compartmentation in which the mass isotopic distributions of malate may differ according to its subcellular localisation.

In *Leishmania spp.* malate serves as both an intermediate of the TCA cycle, within the mitochondria, and as an intermediate in PEP fermentation to succinate within the glycosome. Indeed several intermediates (fumarate, OAA, and succinate) are located in both the glycosome and mitochondria. Indeed OAA and malate may also be found in the cytosol. This compartmentalisation adds another level of complexity to the analysis of eukaryotic cells that is not found in prokaryotes. Unfortunately, currently available techniques are unable to measure compartment-specific metabolite labelling patterns. Commonly, the existence of compartments is ignored in ^13^C-based metabolic flux modelling by assuming uniform labelling patterns of individual metabolites inside multiple compartments. No studies currently exist to prove or disprove this assumption [50]. Indeed, the analyses presented in this work have only been made feasible with the recent development of the novel experimental methods for the detection of labelling patterns inside free intracellular metabolites. It is expected that within the next decade, further advances in the experimental and measurement techniques will allow compartment specific metabolite measurements, and this level new resolution will make ^13^C-based metabolic approaches even more powerful.

Due to potential reversibility of all reactions involved in topping up the OAA and malate nodes, no definitive conclusions can be drawn about the net flux values through the TCA cycle and GSF pathway. However in the future, if enough metabolic network nodes are analysed and the extracellular fluxes are measured, the resulting information can be assembled to compute net fluxes. Moreover, future studies into enzyme reversibility using approaches such as those recently developed by Kleijn and co-workers [51], could be incorporated into this model. Nevertheless, the results for both nodes point to relatively high activity in the TCA cycle, which is also supported by the high production of carbon dioxide reported earlier [34]. Therefore, while some of the relative flux differences can be explained by reaction reversibility, the flux through the TCA cycle might still be higher than that through the GSF pathway. Hence, while the flux through the TCA cycle is not expected to be eight times the value of the GSF flux, as this would interfere with the energetics inside the glycosome, it could still be relatively higher. Its currently believed that approximately half of the PEP produced by glycolysis is imported back into the glycosome for fermentation to succinate in order to regenerate the investment of ATP and NADPH made in the preparatory stages of glycolysis. This would severely restrict the availability of PEP for import into the TCA cycle as OAA (f_15_) [34].

In summary, methodologies developed in this work contribute to the rapid estimation of *in vivo* fractional fluxes (or flux ratios) in *L. mexicana* parasites, based on GC-MS ^13^C-based metabolomics measurements of free metabolic intermediates. Ultimately, ^13^C-based flux modelling provides a high resolution on key areas of metabolism, and the approach presented here and its further development will result in a comprehensive tool for the elucidation of molecular pathogenicity, and precise measurements of the parasite’s metabolic response to different perturbations of interest, such as genetic modifications, nutritional and other environmental changes, including, the effect of the existing and new candidate drugs, and in different life cycle stages. It is hoped that due to the generality of the problem formulation, the approach will also find a wider application to ^13^C-MFRA studies in various other organisms.

## Acknowledgments

The authors would like to acknowledge Metabolomics Australia and David de Souza for the use of their GC-MS in the collection of the *Leishmania* data sets.

